# Prolonged impact of fire on peatland fungi despite rapid recovery of vegetation, prokaryotes, and soil physicochemistry

**DOI:** 10.64898/2026.02.20.707020

**Authors:** Layla Maas, Erik Verbruggen, Marco Cosme, Tobias Ceulemans, Steven Jacobs, Yvonne Liczner, Keunbae Kim, Karen Vancampenhout, Rudy van Diggelen, Willem-Jan Emsens

## Abstract

Climate change is increasing the frequency of wildfires in ecosystems that historically rarely burn, such as wet heaths and peatlands, thereby threatening carbon storage, biodiversity, and ecosystem functioning. We conducted a three-year, multi-level study to assess early post-fire recovery trajectories of soil physicochemical properties, vegetation, and soil microbial communities in a wet peatland–heathland mosaic affected by a flaming wildfire. Using a paired-plot design of burned and adjacent intact plots, we observed immediate spikes in bioavailable nitrogen (NH₄⁺, NO₃^−^) and phosphorus (P_Olsen_) and a reduction in soil moisture in burned plots, yet two years later these parameters had normalized, indicating rapid abiotic recovery. Vegetation was also strongly altered in the year of the fire, quantifiable by a distinct destruction of herb, moss, tree and litter cover. Although initial regrowth was dominated by a relatively fast resprouting of the graminoid *Molinia caerulea*, its absolute cover in burned plots never exceeded its cover in intact plots, suggesting this species did not expand post-fire. More typical peatland and wet heath species, including ericoid shrubs and *Sphagnum* mosses, recovered more gradually but largely returned to pre-fire levels within the timespan of our study, highlighting high vegetation resilience.

Soil microbial communities showed contrasting responses. Prokaryotic communities shifted immediately after burning but largely recovered within one year. Fungal communities, however, exhibited stronger and more persistent changes and followed a distinct recovery trajectory shaped by succession of immediate and delayed fungal responders. Overall, pyrophilous and fire-tolerant fungi, such as *Coniochaeta* spp., increased, as did many presumably generalist or opportunistic saprotrophs. Litter and wood-associated saprotrophs as well as many mycorrhizal taxa, however, declined. Ongoing fungal shifts occurred even after soil chemistry and vegetation had largely returned to baseline, reflecting a temporary decoupling between above- and belowground communities that may have cascading effects on ecosystem functioning. In conclusion, our results reveal differential recovery trajectories across the soil–microbiome–vegetation interface and highlight that seemingly rapid abiotic and aboveground biotic recovery can mask prolonged microbial disruptions. We emphasize the importance of multi-level assessments for understanding ecosystem resilience.

**Highlights:**

- Soil physicochemistry, vegetation and prokaryotes recovered rapidly after a peatland wildfire
- Fungal communities lagged behind and followed a slower recovery trajectory
- The timing and duration of fungal responses to fire varied across taxa and included immediate or delayed as well as short-lived or persistent responders
- There was a mismatch between vegetation and fungal recovery trajectories, evidenced by a transient post-disturbance decoupling between above- and belowground biotic communities
- Presumed aboveground recovery can mask prolonged belowground disruptions, with potential implications for decomposition, nutrient cycling, and plant–microbe interactions

## 1. Introduction

Wildfires are key ecological disturbances that shape biogeochemical cycling and biotic dynamics across many ecosystems (Caiafa et al., 2023; Flanagan et al., 2020). However, climate change is increasing fire frequency, intensity, and severity beyond historical regimes (Cunningham et al., 2024), exposing ecosystems to novel or intensified disturbance pressures (Adkins et al., 2020; K. Nelson et al., 2021; Enright et al., 2022). Understanding how affected systems respond and recover to wildfires is essential to deepen our understanding of ecosystem resilience as well as to predict future ecosystem functioning.

Peatlands are among the most critical ecosystems in this context. Sustained water saturation and anoxic conditions allow them to accumulate vast amounts of organic carbon, estimated at ∼500 Gt globally (Gorham, 1991; Yu, 2012). Historically, natural wildfire occurrence in many temperate peatlands has been very low to nonexistent, but widespread drainage, land-use change, and climate-driven drought increasingly render them flammable (Turetsky et al., 2014; Rein & Huang, 2021). Fires in these systems therefore pose a disproportionate risk to long-term carbon storage and to their specialized biodiversity (Belova et al., 2014; Davies et al., 2023; Kelly et al., 2023; Marfella et al., 2025). Yet, in contrast to fire-adapted ecosystems such as savannas, boreal or Mediterranean forests, post-fire recovery processes in peatlands remain poorly resolved and underexamined.

Across ecosystems, the immediate effects of fire commonly include combustion of vegetation and litter, deposition of ash, and subsequent increases in soil pH and nutrient availability (Ferrenberg et al., 2013; Certini et al., 2021). Vegetation responses depend strongly on life-history strategies: species with protected belowground organs, such as many graminoids, generally resprout rapidly, whereas bryophytes and lichens often rely on recolonization from unburned refugia (Bouskill et al., 2022; Davies et al., 2023). When post-fire environments favor a subset of tolerant or opportunistic plants, persistent shifts in community composition may occur, potentially altering successional trajectories and even alter future fire regimes (Davies et al., 2023).

In contrast to vegetation, belowground responses are generally less well understood. Fire can directly reduce microbial biomass through direct heat damage and mortality (Hart et al., 2005; Pérez-Valera et al., 2019; Certini et al., 2021), or it can indirectly shape the microbiome by transforming the biogeochemical environment via direct host mortality, altered nutrient pools, ash inputs, and accumulation of pyrogenic organic matter (A. R. Nelson et al., 2022; Bouskill et al., 2022; Allingham et al., 2024). These changes may ultimately modify competitive interactions and favor fast-growing as well as stress- or heat-tolerant taxa (Ferrenberg et al., 2013; Dove et al., 2022; A. R. Nelson et al., 2022). Numerous studies, for example, report that bacterial communities tend to recover faster than fungi, frequently resulting in increased bacterial-to-fungal ratios after fires (Certini et al., 2021; Pulido-Chavez et al., 2023; Hopkins et al., 2025).

Such compositional shifts may cascade to ecosystem functioning because soil microbes regulate pivotal ecosystem processes such as decomposition, nutrient mobilization, and plant nutrient acquisition. A hypothetical reduction in the abundance of mycorrhizal fungi, for instance, could constrain plant re-establishment, whereas proliferation of opportunistic saprotrophs may accelerate turnover of newly available organic matter (Sun et al., 2015; Whitman et al., 2019; Certini et al., 2021). Reported post-fire recovery times vary widely between studies, strongly dependent on ecosystem type and fire severity (Hu et al., 2023; Hopkins et al., 2025), ranging from rapid succession within a year (Joukhajian et al., 2026) to persistent differences decades after fire (Pérez-Valera et al., 2017; Pressler et al., 2019; Dove et al., 2022), the latter suggesting a catastrophic regime shift.

Despite the growing body of work on fire-affected ecosystems, two major gaps remain. First, most wildfire studies focus on either soil, vegetation or microbes, while integrated assessments of the joint recovery trajectories of all three actors combined over several consecutive years are, to our knowledge, very rare (but see e.g. Joukhajian et al., 2026). Consequently, we know little about whether above- and belowground communities reassemble synchronously or become temporarily decoupled after a fire event. Second, ecosystems such as forests and grasslands are generally over-represented in fire research (Hopkins et al., 2025), while studies on peatland fires are more scarce, even though peatlands lack evolutionary adaptation to frequent burning and may therefore respond differently from, and more profoundly then, naturally fire-prone systems.

Here, we present a three-year, multi-level analysis of post-fire recovery in a temperate peatland–heathland mosaic affected by a flaming wildfire in northern Belgium. Using paired burned and intact plots, we examined (1) how fire altered soil physicochemical properties, (2) how vegetation, fungal, and prokaryotic communities responded and changed through time, and (3) whether recovery trajectories of above- and belowground communities were synchronized or decoupled. By combining plant, microbial and soil perspectives, our study provides one of the first integrated evaluations of ecosystem resilience and community reassembly in a burned down peatland-heathland system.

## 2. Materials and methods

### 2.1 Study site

The study was conducted in nature reserve “Landschap De Liereman” in Oud-Turnhout (51.338186, 5.027168), Belgium, where an uncontrolled wildfire affected approximately 30 hectares on April 22^nd^ 2020. The fire event was fierce and high-intensity and appeared catastrophic, but it was also rapid and seemingly shallow: although aboveground biomass was charred across the affected area, the flaming fire did not evolve into a smouldering burn and was extinguished within one day. We believe its short duration was due to the fire naturally running out of fuel (litter) in combination with extinguishing efforts by the fire department. The burned area lies within a gently sloping peatland complex representing the upper (source) section of a lateral percolation mire system. The site features a heterogeneous mosaic of nutrient-poor habitat types, forming a gradient from acidic moist heath to weakly buffered poor fen habitat and remnant bog communities. Key vegetation types include wet heath (dominated by *Erica tetralix* L.), locally transitioning into moist *Calluna vulgaris* (L.) Hull heaths, and more degraded communities dominated by the graminoid *Molinia caerulea* (L.) Moench. Transitional poor fen vegetation with *Narthecium ossifragum* (L.) Huds. is widespread, reflecting early successional stages toward raised or transitional bog. Additional affected habitats include *Myrica gale* L. vegetation and relict bog fragments with active *Sphagnum* carpets and *Vaccinium oxycoccos* L.

### 2.2 Site selection, study design and sampling

Immediately following the fire event, the boundaries of the burned area were clearly visible with most of the aboveground biomass absent or charred, allowing for an accurate delineation of burned and unburned intact zones. During a subsequent field survey, 15 sub-locations were selected across the affected landscape. At each sub-location, two paired permanent plots (2 × 2 m) were established: one located within a burned area (henceforth referred to as “burned”) and the other in an adjacent intact control area (“intact”). This was possible since the fire had affected the area in a very heterogeneous manner, leaving many subsites untouched. In total, 30 plots were installed (15 pairs) in a paired-plot design. The plots were distributed proportionally across the dominant habitat types within the affected area, ensuring representation of the full moisture and habitat gradient present at the site. The study was conducted over three consecutive growing seasons, beginning immediately in the year of the fire (2020) and continuing through the summer of 2022. Sampling of soil, as well as the making of vegetation relevees, was done each year in the last week of July. Composite soil samples were obtained from four subsamples within each plot: here, the surface layer (0-5 cm) was sampled due to its highest exposure to fire. The composite samples were then stored at -18°C until further processing. Simultaneously, we always collected a second pairwise batch of samples of known volume (using 100 cm^3^ Kopecky rings) for calculation of soil bulk density.

### 2.3 Vegetation surveys

Vegetation surveys were performed annually in all 30 plots: all plant species were identified, and their respective absolute covers (expressed as % of the entire plot) were visually estimated in each plot. In addition, we also calculated relative species covers, which we defined as the cover of the individual species divided by the total cumulative cover of all species (*100). Overall vegetation structure was visually estimated by the total absolute cover (%) of the herb, moss, tree (including tree seedlings and saplings), and litter layers.

### 2.4 Soil physicochemical analyses

Soil subsamples of the first (2020) and last (2022) study year were used for soil physicochemical analyses.

Concentrations of bio-available nitrate (NO_3_ ^−^) and ammonium (NH_4_ ^+^) were determined on fresh soil subsamples by extraction with KCl (2h shaking in 1M KCl; 1:5) and subsequent analysis on an auto-analyzer system (SAN++, Skalar). Soil acidity (pH_KCl_) was determined in the extracts using portable equipment (Hannah Instruments, HI99121 with HI12923 electrode).

To determine the acid-neutralizing capacity of the soil, cation exchange capacity (CEC) and base saturation (BS) were determined by adding 50.0 ml of 1M ammonium acetate (brought to pH 7 with NH_3_OH) to 5 g of soil subsample. The mixture was then shaken for 1 h and filtered, after which cations were measured using ICP (iCAP 6300 Duo, Thermo Scientific). Exchangeable H^+^ was determined by titration following (Brown, 1943). Total CEC (in meq 100g^−1^) was calculated as the cumulative sum of all exchangeable cations (Ca^2+^, Mg^2+^, K^+^, Na^+^, Fe^2+^, Al^3+^, Mn^2+^, H^+^), and base saturation (in %) was calculated as the percentage of base cations (Ca^2+,^ Mg^2+^, K^+^, Na^+^) relative to total CEC.

Bioavailable phosphorus (P_Olsen_) was determined by NaHCO_3_ extraction on ground and dried (40°C) soil (Olsen, 1954) and P was measured on an auto-analyzer system (SAN++, Skalar).

Total soil nitrogen (N) and total phosphorus (P) concentrations were quantified through complete digestion in H_2_SO_4_ of ground oven-dried (70°C) soil and subsequent analysis on an auto-analyzer system (SAN++, Skalar).

Soil bulk densities (kg L^−1^) and moisture contents (%) were determined by drying the second set of samples of known volume (100 cm^3^) at 105°C, after which organic matter contents (%) were determined gravimetrically by loss-on-ignition (4h at 550°C). To calculate element pool sizes (mmol L^−1^) in the soil, we multiplied element concentrations (mmol kg^−1^) with bulk densities of the corresponding samples.

### 2.5 Microbial analyses

Microbial communities were analysed with the use of DNA metabarcoding. Approx. 5 ml of frozen soil subsamples from the years 2020, 2021 and 2022 were lyophelized, fragmented using scissors, subsampled to 1 ml and then pulverized in a ball-mill to obtain a representative sample. Approx. 0.15 g was then subjected to DNA isolation using the Powersoil Pro kit (Qiagen, Venlo, the Netherlands). The fungal ITS1 region was amplified using general fungal primers ITS1f (Gardes & Bruns, 1993) and ITS2 (White et al., 1990). For prokaryotes, an approximately 250 bp stretch within the V4 region of the 16S rRNA was amplified with the F515/R806 primer set (Caporaso et al., 2011). Each primer contained Illumina adapters at the 5′ ends to facilitate adding barcodes and sequencing adapters in a second PCR (“Nextera” procedure). Each 25 μl reaction mixture contained 2 μl of template, 16.3 μl of H2O, 0.2 μM of each forward and reverse primer, 5 μl PCR buffer, 0.5 μl dNTPs and 0.2 μl Phusion High-Fidelity DNA polymerase (New England Biolabs, Ipswich, MA, USA). PCR conditions were as follows: initial denaturation at 98 ◦C for 60 s, followed by 30 (fungi) or 22 (prokaryotes) cycles of denaturation at 98 ◦C for 30 s, annealing at 55 ◦C for 30 s, extension at 72 ◦C for 30 s; and an additional extension of 72 ◦C for 10 min. PCR products were run on a 1.5% agarose gel to confirm successful PCR amplification and to confirm that negative controls were empty (and in case of failure the procedure was repeated). Successful PCR products were diluted 50 x and a second PCR was performed using dual barcoded primers with Illumina adapters in an identical PCR mix as above except using 3 μl of template and adjusting the water content accordingly. The conditions were: 98 ◦C for 60 s, 12 cycles: at 98 ◦C for 10 s, 63 ◦C for 30 s, 72

◦ C for 30 s; and 72 ◦C for 5 min. PCR products were run on an agarose gel and successful amplicons were pooled into a single library. The library was purified using Machery Nagel gel purification kits QIAquick Gel Extraction Kit (Qiagen, Venlo, the Netherlands) and quantified using qPCR (KAPA Library Quantification Kits, Kapa Biosystems, Wilmington, MA, USA). The sequencing was performed using the Illumina MiSeq platform (Illumina Inc; San Diego, CA, USA) with 2×300 cycles for forward and reverse reads. Microbial Amplicon Sequence Variants (ASVs) were defined following the Dada2 pipeline (Callahan et al., 2016). The resulting ASVs were annotated using the SILVA database v138.1 and UNITE v9.0 (Abarenkov et al., 2023) for 16S and ITS, respectively, and non-target sequences removed.

### 2.6 Data analyses

All statistical analyses were conducted in R version 4.3.3 (R core team, 2025) and significance for statistical tests was accepted at P < 0.05.

To account for uneven sequencing depth of ASV reads between samples, rarefaction was performed to a sequencing depth of 10,000 sequences/sample for fungi and 5,000 sequences/sample for prokaryotes with the rrarefy() function from vegan (version 2.6.4) (Oksanen et al., 2022). After rarefaction, 3,203 fungal and 8,164 prokaryotic ASVs were left. Of the fungal samples, 5 did not reach the sampling depth with a range of 2,010 to 8,615 reads. For prokaryotic samples, 6 samples did not reach sampling depth with a range of 3,189 to 4,980 reads. Samples that did not reach sufficient reads were normalised. The effect of fire on community composition was assessed with a nonparametric multivariate ANOVA (PERMANOVA) (Anderson, 2001) with the adonis2 function from vegan based on bray-curtis dissimilarities. Prior to distance calculation, ASV counts and vegetation cover data were square root transformed. Non-metric multidimentional scaling (NMDS) was used for ordination of vegetation and microbial communities, which was performed with the function metaMDS() from vegan. Differential abundance analysis between burned and intact (control) plots of microbial taxa was carried out using DESeq2 (Love et al., 2014) on raw count data (before rarefaction) in order to include rare taxa. The threshold for significance was set at log₂ fold change > |2| and an adjusted p-value of 0.05. ASVs with a significant change in relative abundance are considered responders. Responders are divided in ASVs that responded in the first year (immediate), and those that responded in 2021 or 2022 (delayed), as well as those with an increase in relative abundance (log₂ fold change > 2; positive) or a decrease in relative abundance (log₂ fold change > -2; negative), resulting in four categories of responders (1) Immediate positive, (2) Immediate negative, (3) Delayed positive, (4) Delayed negative.

Fungal lifestyles were assigned with FungalTrait database (Põlme et al., 2020). Significant differences in fungal lifestyles and soil physicochemical properties after fire were tested using paired t-tests or Wilcoxon signed-rank tests depending on the normality of the paired differences, determined with the Shapiro-Wilk test. All tests were performed separately for each year in order to assess changes over time. All visualisations were created with the use of ggplot2 (Wickham, 2016).

Correlation between prokaryotic and fungal community composition with vegetation community composition was analysed with a mantel test (Spearman’s rank correlation) from vegan.

## 3. Results

### 3.1 Fire-induced changes in soil physicochemical properties

Wildfire did not result in any significant immediate (2020) or delayed (2022) changes in top soil organic matter contents, bulk densities, pH, base saturation or total soil pools of nitrogen (N) or phosphorus (P) (Table 1, Suppl. Fig. S1). There was however a strong and significant increase in bio-available P (P_Olsen_), extractable ammonium (NH_4_ ^+^), and extractable nitrate (NO_3_ ^−^) in the burned plots in the year of the fire (2020), whereas soil moisture contents had slightly decreased (Table 1, Suppl. Table S1). In the third growing season (2022) however, there were no longer any measurable differences in soil nutrient availabilities or moisture contents, but there was a minor but significant increase in soil cation exchange capacity (CEC), which seemed solely attributable to a slight but significant increase in protons (H^+^) in the burned plots (Table 1).

**Table 1:** Comparison of means (± SD) of soil variables between burned and intact plots measured in 2020 and 2022, or only in 2022 (CEC and BS). Depending on assumptions of normality of the paired differences, either paired t-test results (t-statistic) or Wilcoxon-signed-rank (V-statistic) results are reported per soil variable. Significance is indicated as follows: ***p<0.001; **p<0.01; *p<0.05.

| Parameter | Unit | Year | Mean ± SD<br>Intact (n = 15) | Mean ± SD<br>Burned (n = 15) | t-statistic<br>df = 14 | V-statistic |
| --- | --- | --- | --- | --- | --- | --- |
| Bulk density | kg L<sup>-1</sup> | 2020 | 0.3 ± 0.4 | 0.3 ± 0.4 | | 71 |
| | | 2022 | 0.3 ± 0.4 | 0.3 ± 0.3 | | 56 |
| Moisture content | % | 2020 | **74.2 ± 25.6** | **68.4 ± 26.7** |  | **23 \*** |
| | | 2022 | 72.4 ± 25.7 | 69.1 ± 21.6 | -1.2 | |
| $\text{NH}_4^+$ | $\mu\text{mol L}^{-1}$ | 2020 | **1151.7 ± 1383.0** | **3843.9 ± 2636.3** | **3.5\*\*** | |
| | | 2022 | 452.7 ± 435.6 | 552.0 ± 847.7 | | 57 |
| $\text{NO}_3^-$ | $\mu\text{mol L}^{-1}$ | 2020 | **115.1 ± 255.4** | **192.4 ± 241.2** | **2.1\*** | |
| | | 2022 | 48.7 ± 92.7 | 49.9 ± 124.9 | | 37 |
| Total N | mmol L<sup>-1</sup> | 2020 | 258.1 ± 409.9 | 286.8 ± 425.2 | | 58 |
| | | 2022 | 162.8 ± 81.0 | 227.7 ± 128.9 | 1.71 | |
| Organic matter | % | 2020 | 71.1 ± 31.9 | 69.5 ± 32.9 | | 68 |
| | | 2022 | 73.5 ± 30.7 | 72.8 ± 28.7 | | 64 |
| Total P | mmol L<sup>-1</sup> | 2020 | 4.1 ± 5.1 | 5.5 ± 7.9 | | 114 |
| | | 2022 | 2.5 ± 1.1 | 3.0 ± 1.7 | 1.11 | |
| $P_{\text{Olsen}}$ | $\mu\text{mol L}^{-1}$ | 2020 | **73.6 ± 34.9** | **225.0 ± 190.9** | | **114\*\*\*** |
| | | 2022 | 69.8 ± 34.9 | 125.9 ± 174.5 | | 83 |
| pH<sub>KCl</sub> | | 2020 | 3.3 ± 0.2 | 3.5 ± 0.2 | 1.44 | |
| | | 2022 | 3.5 ± 0.3 | 3.5 ± 0.2 | -0.79 | |
| CEC | meq 100g<sup>-1</sup> | 2022 | **44.1 ± 20.2** | **51.3 ± 20.4** |  | **87\*** |
| Base saturation (BS) | % | 2022 | 21.2 ± 8.8 | 18.1 ± 7.2 | -1.45 | |
| $\text{Al}_{\text{exch}}$ | meq 100g<sup>-1</sup> | 2022 | 0.3 ± 0.2 | 0.2 ± 0.1 | | 28 |
| $\text{Ca}_{\text{exch}}$ | meq 100g<sup>-1</sup> | 2022 | 5.9 ± 3.4 | 5.5 ± 3.2 | -0.35 | |
| $\text{Fe}_{\text{exch}}$ | meq 100g<sup>-1</sup> | 2022 | 0.2 ± 0.2 | 0.3 ± 0.1 | | 43 |
| $\text{H}_{\text{exch}}$ | meq 100g<sup>-1</sup> | 2022 | **33.5 ± 14.6** | **40.9 ± 16.3** | | **102\*** |
| $\text{K}_{\text{exch}}$ | meq 100g<sup>-1</sup> | 2022 | 1.1 ± 0.9 | 0.9 ± 0.6 | -1.16 | |
| $\text{Mg}_{\text{exch}}$ | meq 100g<sup>-1</sup> | 2022 | 2.0 ± 1.3 | 2.1 ± 1.3 | 0.051 | |
| $\text{Mn}_{\text{exch}}$ | meq 100g<sup>-1</sup> | 2022 | 0.1 ± 0.1 | 0.1 ± 0.1 | | 46 |
| $\text{Na}_{\text{exch}}$ | meq 100g<sup>-1</sup> | 2022 | 1.28 ± 1.0 | 1.3 ± 0.9 | -0.24 | |

### 3.2 Fire-induced changes in overall biotic community compositions

Over the course of three growing seasons, we counted a total of 27 vascular plant taxa and 8 *Sphagnum* species, as well as a total of 3,294 unique fungal and 8,660 prokaryotic ASVs.

NMDS ordinations indicated immediate fire-induced shifts in community compositions of biota, with PERMANOVA results confirming significant compositional changes across vegetation (F(1,29) = 2.96, p = 0.009), prokaryotic (F(1,27) = 1.50, p = 0.046), and fungal (F(1,29) = 3.10, p < 0.001) communities in the year of the fire (2020; Fig 1a-c). Vegetation and prokaryotic communities, however, exhibited relatively minor ordination shifts and rapidly converged with intact communities, with no more significant differences detected by 2021. In contrast, fungal communities displayed stronger immediate fire-induced shifts. Three years later, fungal community composition in burned plots still remained marginally different from intact plots (F(1,28) = 1.37, p = 0.053), despite a visual trend of gradual convergence (2022; Fig 1c). Together, these patterns indicate a rapid post-fire recovery of vegetation and soil prokaryotes, but a delayed recovery of soil fungi.

**Fig. 1:**
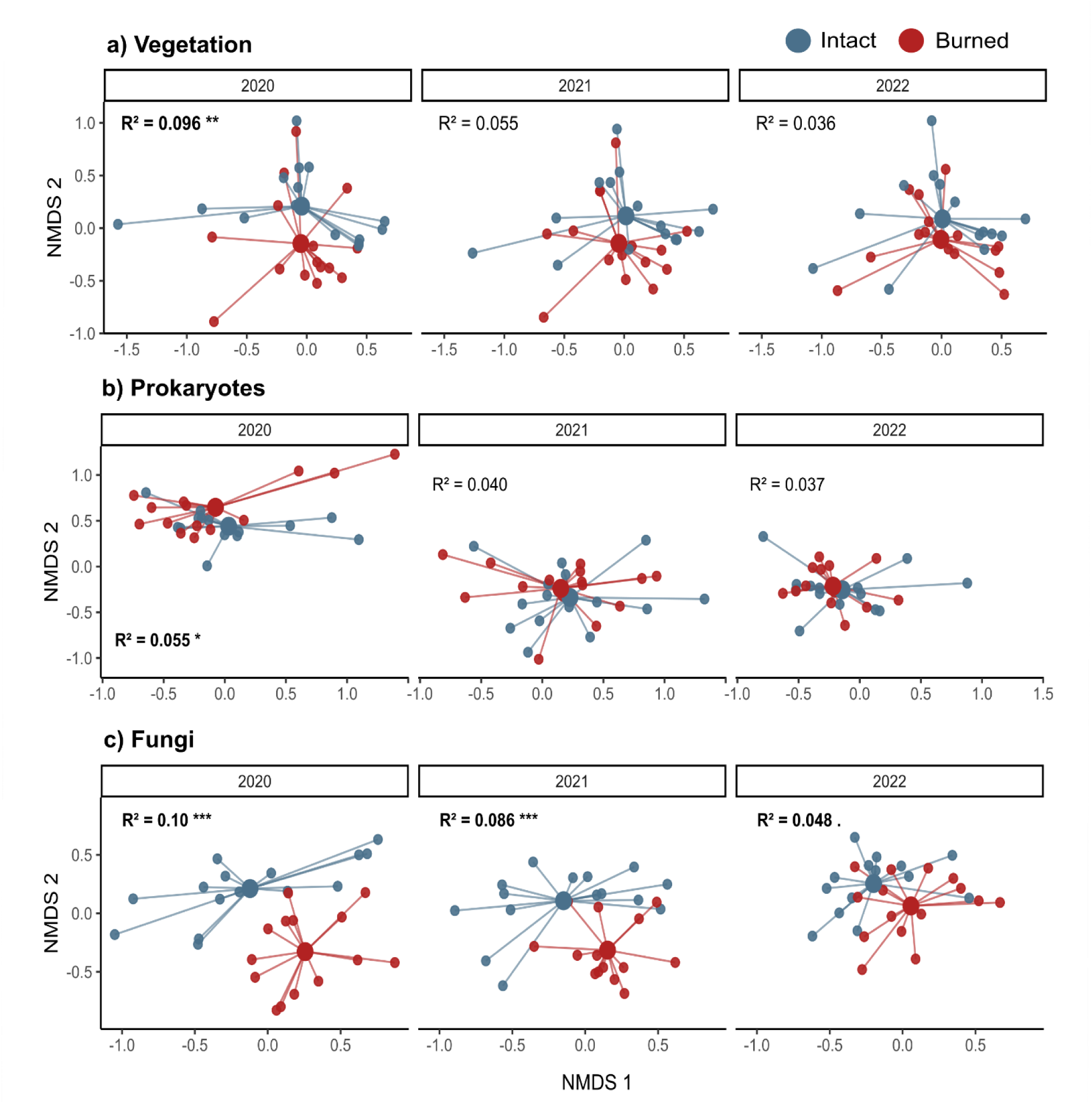
Ordination (non-metric multidimensional scaling; NMDS) of post-fire (a) vegetation (K = 3, stress = 0.13), (b) prokaryotic (K = 3, stress = 0.13), and (c) fungal community composition (K = 3, Stress = 0.17) in burned and intact plots for the period 2020-2022. The analyses are based on Bray-Curtis dissimilarity indices of the sample*species and sample*ASV matrices. Every point represents community composition in a unique plot; centroids represent the average of all samples. The PERMANOVA R^2^ is given for the effect of burning, with statistical significance indicated as follows: ***p<0.001; **p<0.01; *p<0.05.

### 3.3 Post-fire recovery of vegetation structure and individual plant species

The fire led to a strong reduction in litter cover, which remained lower in the burned plots as compared to intact plots for three consecutive years (2020: t(14) = -3.85, p =0.002; 2021: t(14) = -4.38, p=0.001, 2022: t(14) = -4.22, p=0.001) (Fig. 2a). In addition, we measured a significant reduction in herb cover in the first and second years post-fire (t(14) = -7.75, p < 0.001 and t(14) = -3.87, p= 0.002) (Fig. 2b), but there was convergence in year three (W = 21, p= 0.051). Total moss cover, consisting of *Sphagnum* spec., was only significantly lower in the year of the fire (t(14) =-2.65, p= 0.019), displaying a swift recovery trend (Fig. 2c). The tree layer, including seedlings and saplings, shows a similar trend to herbs and mosses but the difference between burned and intact plots was only slightly significant in 2022 (V=2.5, p=0.033) (Fig. 2d). Immediate vegetation recovery was dominated by rapid regrowth of the graminoid *Molinia caerulea*, the relative cover (= individual species cover / total cumulative cover of all species) of which was higher in the burned plots (Suppl. Fig. S2). This pattern of relative *Molinia*-dominance remained throughout all years (2020: t(14)=3.61, p = 0.002; 2021: t(14)=4.40, p=0.001; 2022: t(14)=4.22, p=0.001). However, the absolute cover of *Molinia* in burned plots never significantly surpassed that of *Molinia* in intact plots (Suppl. Fig. S3, 2020: t(14)= -1.59, p=0.133; 2021: t(14)=0.89, p=0.386; 2022: t(14)=0.37, p=0.715). Other plant species, including key species such as the ericoid shrubs *Calluna vulgaris* and *Erica tetralix* as well as individual species of *Sphagnum,* exhibited a slightly more gradual recovery pattern over the years (Suppl. Figs S2 and S3).

**Fig 2:**
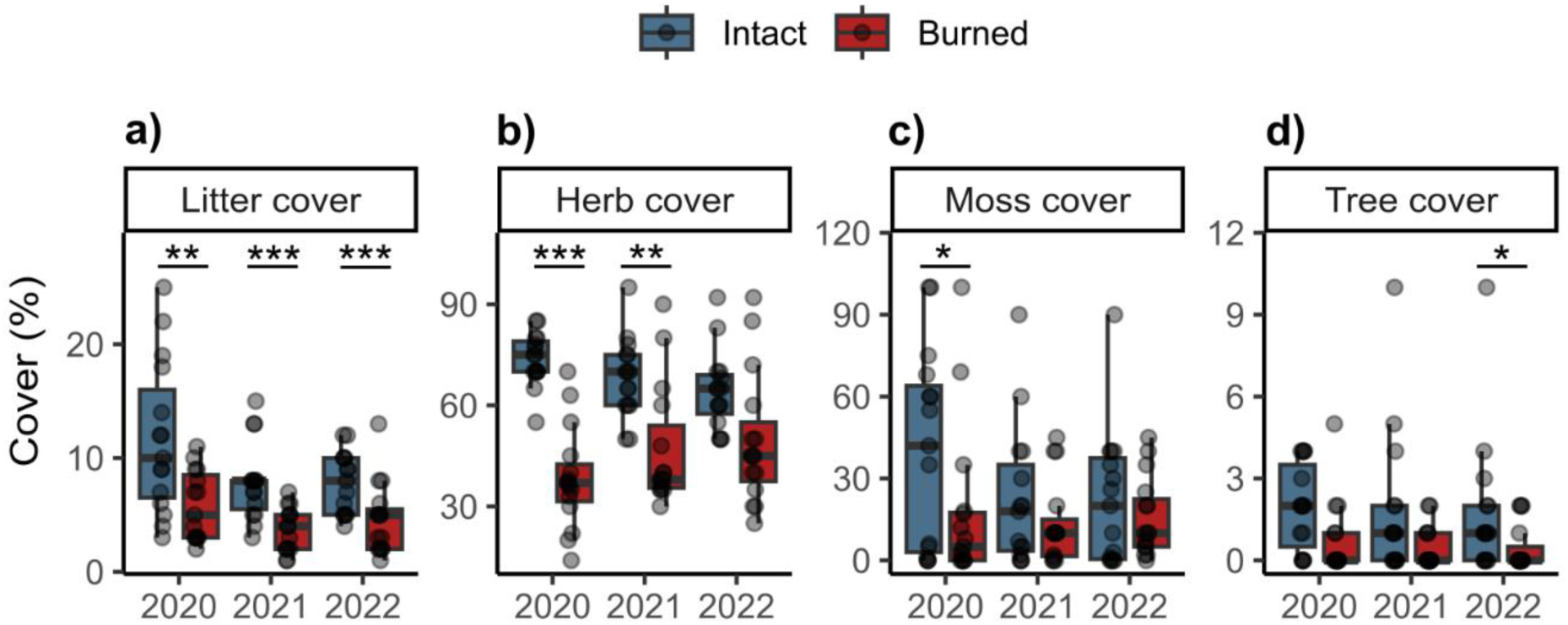
Absolute cover of (a) litter, (b) herbs, (c) mosses and (d) trees (including seedlings and saplings) in burned (n=15) and intact (n=15) plots from 2020 to 2022. Significance is indicated as follows: ***p<0.001; **p<0.01; *p<0.05.

### 3.4 Post-fire responses of microbial taxa

A wide range of prokaryotic and fungal taxa showed significant changes in relative abundance following the fire (log₂ fold change > |2|, adjusted p < 0.05), with the largest number of differentially abundant ASVs observed in the year of the fire (2020) and the fewest in the final post-fire sampling year (2022; Fig. 3).

**Fig. 3:**
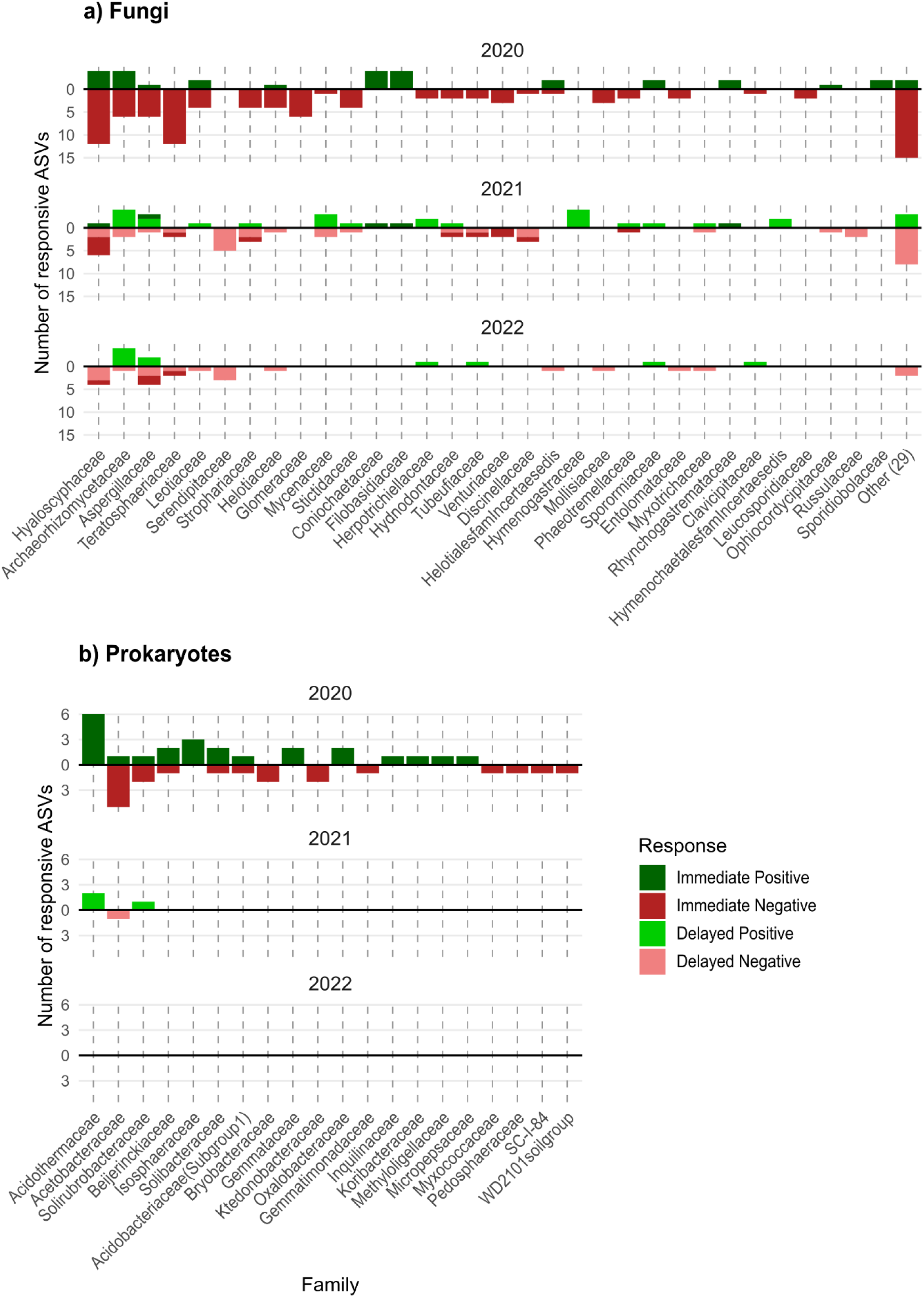
Responses of microbial ASVs grouped by family with a significant change in relative abundance (Log₂ Fold change > |2|, p < 0.05) following fire as identified by DESeq. This change can either be an increase (Log₂FC > 2; green) or decrease (Log₂FC < -2; red) in relative abundance after fire. Responses are grouped by those that show a significant change in the first year (“Immediate response”) and those that only react in 2021 or 2022 (“Delayed response”). Fungal families with only one responsive ASV are grouped and marked as ‘other’ (n=29). ASVs with unknown family classification are not shown in this figure (but see Suppl. Figs S4 & S5).

For fungi, 168 of 1,892 ASVs (9%) were significant responders in 2020, 107 of 1,555 ASVs (7%) in 2021, and 49 of 1,662 ASVs in 2022 (3%), respectively accounting for an average of 36.0%, 17.3%, and 8.7% of the total fungal reads (Fig. 3a). Responses included immediate changes in 2020, delayed responses detected only in 2021 or 2022, and prolonged responses persisting across multiple years (of which 19 ASVs in both 2020 and 2021, 7 in 2021 and 2022, 5 in 2020 and 2022).

Fire-responsive fungi were taxonomically widespread and were dominated by declines, i.e. the majority of fungal responses were negative (Fig. 3a, Suppl. Fig. S5). Clear and unilateral positive responses were largely restricted to a few putative pyrophilous or opportunistic saprotrophic taxa, such as the mold- or yeast-like Coniochaetaceae (predominantly *Coniochaeta cipronana*) and Filobasidiaceae families. Predominantly negative responders were most common among families that contain many mycorrhizal fungi, including Helotiaceae (containing ericoid mycorrhiza), Hyaloscyphaceae (e.g. the ericoid mycorrhizal *Hyaloscypha variabilis)* or Glomeraceae (arbuscular mycorrhiza), but also, for example, in the non-mycorrhizal families Teratosphaeriaceae and Strophariaceae. The majority of fungal families, however, contained ASVs with mixed responses (Fig 3a, Suppl. Fig. S5).

In comparison with fungi, very few prokaryotic taxa were significant responders to the fire (log₂ fold change > |2|, adjusted p < 0.05), with 67 of 7,638 ASVs (0.9%) in 2020, 4 of 1,736 ASVs (0.2%) in 2021, and none of 1,491 ASVs in 2022 (Fig. 3b, Suppl. Fig. S4). In the first two years, these responsive ASVs represented only an average of 5.87% and 2.02% of total prokaryotic reads, respectively. Hence, the majority of responsive ASVs exhibited immediate changes in 2020, and with little or no changes or responses in 2021 or 2022. Also, no ASVs were identified as responders for over more than one year, indicating that changes in prokaryotic relative abundance were short-lived and did not persist beyond a single year. Responsive prokaryotic ASVs were taxonomically widespread and responses were variable, but clear and unilateral positive responders were primarily members of Acidothermaceae, Isosphaeraceae, Gemmataceae and Oxalobacteraceae such as *Massilia sp.,* while negative responders were, amongst others, found in Acetobacteraceae, Bryobacteraceae, and Ktedonobacteraceae (Fig. 3b, Suppl. Fig. S4).

### 3.5 Changes in putative fungal lifestyles

Fungal lifestyles that were significantly more prevalent after fire included general saprotrophs (unspecified) (2020; V=8, p=0.003, Fig. 4a), which was also the most dominant assigned functional group in terms of relative abundance, as well as mycoparasites (2021; t(14)= -3.54, p=0.003, 2022; t(14)=-2.72, p=0.017), Fig. 4f), animal parasites (2022: V=9, p=0.021, Fig. 4e) and dung saprotrophs (2021; V=5, p=0.003, Fig. 4d). Litter saprotrophs (2021; t(14)=2.48, p = 0.027, Fig. 4b) and wood saprotrophs (2020; V=105, p=0.011, 4c) tended to decrease post-fire. Lichenized fungi decreased in the first year (2020: V=45, p = 0.009), but increased in the years after (2021: V=10, p = 0.025; 2022: V=0, p=0.022, Fig. 4g).

**Fig. 4:**
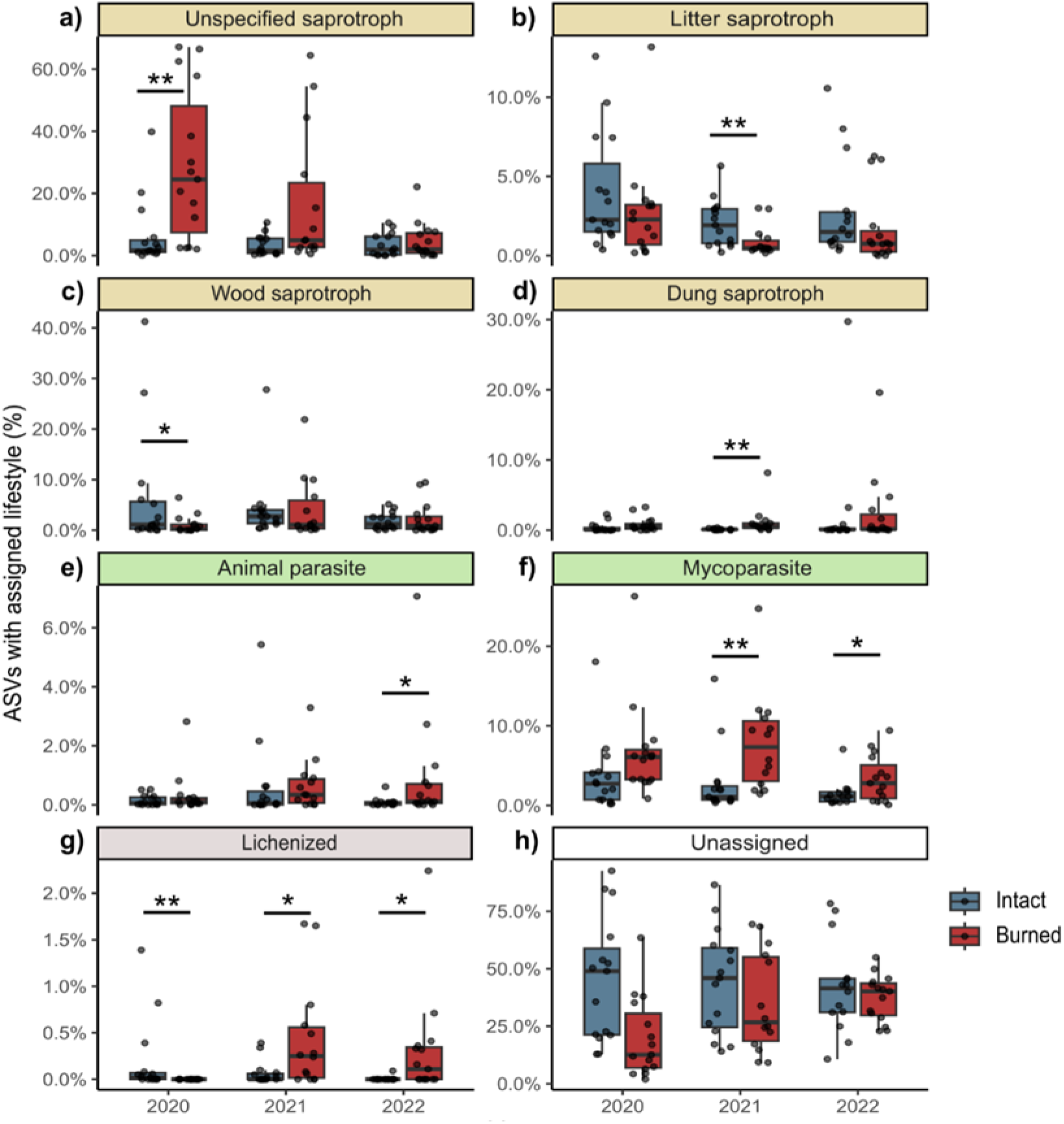
Putative fungal lifestyles in burned and intact plots over time (2020-2022). Percentages represent the proportion of ASVs in samples of burned (n=15) and intact (n=15) plots in a given year that were assigned to a particular fungal lifestyle using the Fungaltrait database. Significance is indicated as follows: ***p<0.001; **p<0.01; *p<0.05.

### 3.6 Fire-induced disruption of plant-fungal relationships

There was a relatively strong correlation between fungal and vegetation community composition in the intact plots across years, with significant positive correlations in 2020 (Mantel rho = 0.478, p < 0.001) and 2021 (Mantel rho = 0.448, p = 0.001) and a marginally significant positive correlation in 2022 (Mantel rho = 0.221, P = 0.074) (Fig. 5a-c).

**Fig. 5:**
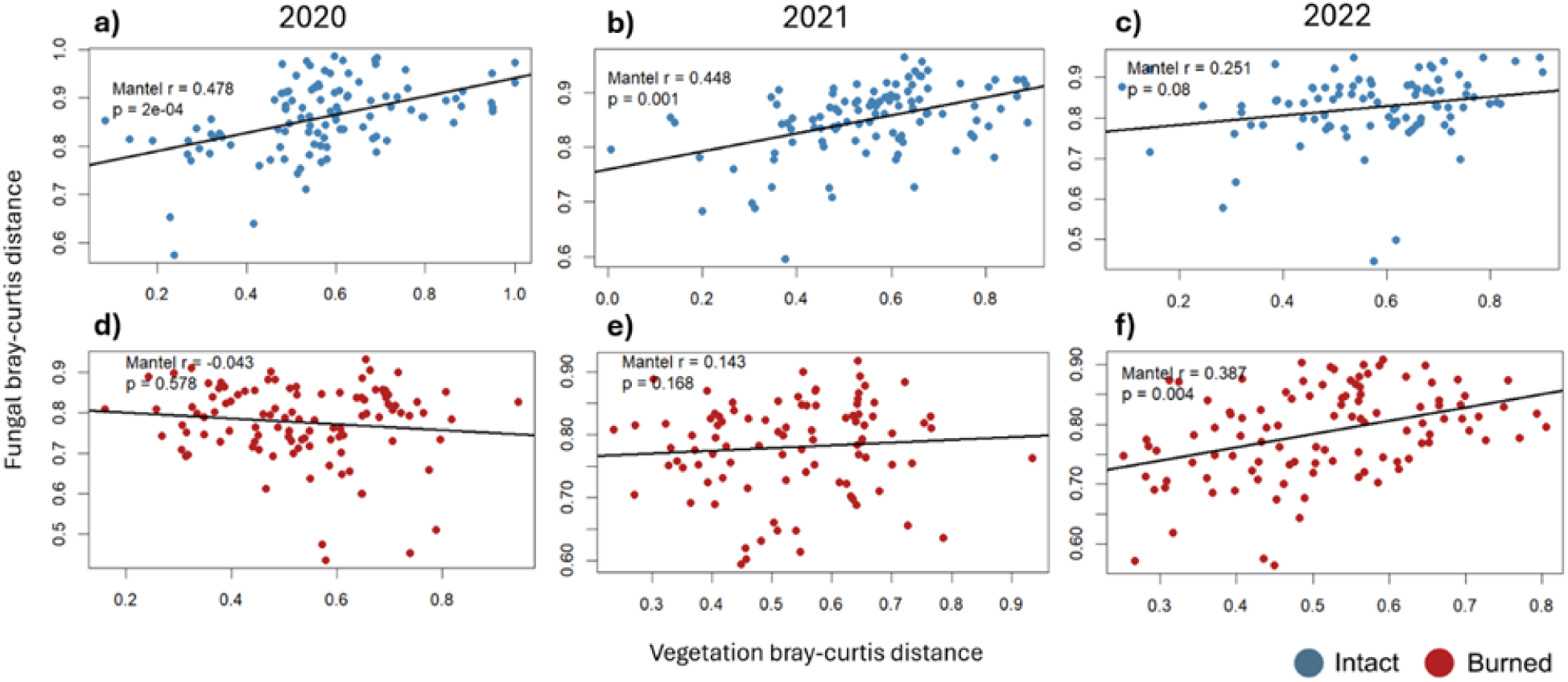
Correlations between fungal and vegetation community composition in intact (panels a-c, n=15) and burned plots (panels d-f, n=15) for the period 2020-2022. Mantel’s correlation coefficients (Spearman’s rho) and p values are indicated on the graphs.

In the burned plots however, fungal community composition was disconnected from vegetation community composition in the first two years after the fire (Mantel rho = -0.043 and 0.143 respectively, p > 0.05), with positive alignment of both communities only in the third year post-fire (Mantel rho = 0.387, p = 0.004). Prokaryotic community composition was not correlated to vegetation community composition at any point in time in either the burned or intact plots (Suppl. Fig. S6).

## 4. Discussion

We conducted an integrated and multi-year assessment of post-wildfire recovery trajectories of soil physicochemical properties, vegetation, and soil microbial communities in a nutrient-poor temperate peatland–heathland mosaic. Using a design of burned-intact plot pairs, we show that soil chemistry and vegetation recovered rapidly, whereas soil microbial communities, particularly fungi, exhibited more pronounced and persistent shifts. These contrasting recovery trajectories highlight a temporary decoupling between above- and belowground community dynamics following a severe perturbation.

### 4.1 Alterations in soil physiochemistry were limited and transient

Fire-induced alterations in soil physicochemical properties were limited in magnitude and duration, which we link to the short-lived and predominantly surficial nature of the flaming fire event. Total soil organic matter content, bulk density, soil pH, base saturation, and total nitrogen and phosphorus pools did not differ between burned and intact plot pairs, suggesting minimal structural degradation of the peat. This is in sharp contrast to findings from studies on smouldering peat combustion, i.e. slow-burning flameless combustion that tends to “disappear” deep into the peatland soil where it can persist for days to weeks and cause biogeochemical havoc (Davies et al., 2013; Rein & Huang, 2021).

The most pronounced edaphic effect in our study was a significant increase in bioavailable phosphorus (P_Olsen_), ammonium (NH_4_ ^+^) and nitrate (NO_3_ ^−^) in the year of the fire, accompanied by a slight reduction in soil moisture content. Such pulses in inorganic nutrients are commonly seen after peat fires (Orlova et al., 2020; van Beest et al., 2019), and are likely associated with sudden large-scale biomass combustion and plant die-off, as well as with potential fire-induced mineralization of organic matter or litter and subsequent ash deposition (Agbeshie et al., 2022; Smith et al., 2001).

These nutritional shifts, however, were transient and no longer detectable by the third growing season after the fire, suggesting rapid abiotic recovery. Since the affected heathland-peatland ecosystem is naturally poor in macronutrients, we hypothesize that the nutrient pulse was rapidly neutralized by a combination of plant uptake, microbial immobilization and leaching through (ground)water percolation. A minor but significant increase in cation exchange capacity (CEC) in burned plots in the third monitoring year was solely attributable to higher exchangeable proton (H^+^) concentrations rather than changes in base cations, and it was not accompanied by consistent shifts in pH or base saturation or overall nutrient availability, suggesting limited ecological relevance.

### 4.2 Rapid recovery of vegetation community composition

The wildfire caused an immediate and pronounced reduction in aboveground vegetation cover in the affected plots, and it heavily impacted initial vegetation community composition. Studies on post-fire vegetation recovery trajectories of European heathland-peatland landscapes often report a rapid recolonization and subsequent expansion by fast-growing graminoids such as purple moor grass (*Molinia caerulea*) (Brys et al., 2005; Jacquemyn et al., 2005), which, if the root system survives, may take advantage of the immediate post-fire spike in nutrient availability (Brys et al., 2005). Even though *Molinia caerulea* is a native species in the region, its dominance in a peatland-heathland landscape is generally considered unwanted as it indicates degradation and it may push the ecosystem toward a species-poor alternative semi-stable state. In line with literature, we found that *Molinia* indeed resprouted rapidly from surviving belowground organs and temporarily increased in relative dominance, i.e. in comparison to other plant species. However, the absolute cover of *Molinia* in burned plots never exceeded the cover in paired intact plots, indicating that fire did not promote long-term expansion. In other words: *Molinia* resprouted rapidly after the fire, but it did not seem to expand. Typical peatland and wet heathland species, including ericoid shrubs and several *Sphagnum* species, recovered more gradually but had largely returned to pre-fire levels within one to two years. This pattern aligns with previous findings of rapid (∼2 years) post-fire recovery of *Sphagnum* spp. following low-severity burns (Grau-Andrés et al., 2017). Such resilience may be attributed to the inherently high tissue moisture content of *Sphagnum*, which may reduce flammability (Shetler et al., 2008). Consistent with this mechanism, many *Sphagnum* patches in our burned plots were not completely combusted despite clear signs of physiological damage (e.g., discoloration and structural degradation; pers. obs. WJ Emsens).

In accordance, overall vegetation composition of burned plots converged with intact plots already within one year after the fire, indicating a rapid recovery of plant communities despite the magnitude of the initial disturbance. As such, vegetation recovery closely tracked the rapid normalization of soil physicochemical conditions. This fast overall convergence suggests a surprisingly high resilience of the peatland–heathland vegetation to a low-severity flaming fire, while the absence of persistent changes in species cover or community composition suggests that fire did not trigger a shift toward alternative vegetation states. We nonetheless believe that, if there had been a more severe and intense combustion of peat and *Sphagna*, plant-soil recovery trajectories would likely have been slower.

### 4.3 Differential responses of soil microbial communities and temporary decoupling from vegetation

Fire triggered strong immediate shifts in soil microbial communities, aligning with the strong shifts in nutrient availability and vegetation, but the nature and duration of these changes differed markedly between prokaryotes and fungi.

For prokaryotes, very few taxa were identified as significant responders to the fire, and responses were largely short-lived: shifts in relative abundances were most evident immediately after the fire, with roughly equal numbers of positively and negatively responding taxa. Strongest positive responders consisted of taxa that are linked with traits favoring heat tolerance, for example thermophillic *Acidothermus* sp. (Acidothermaceae) (Mohagheghi et al., 1986), or traits favoring rapid growth, for example *Massilia sp*. (Oxalobacteraceae) which has been identified consistently as a positive fire responder in a range of ecosystems (Whitman et al., 2019; Caiafa et al., 2023; Pulido-Chavez et al., 2023; Amirhosseini et al., 2025). One year after the fire however, prokaryotic communities already closely resembled those of intact plots, suggesting a rapid recovery on the community level. This is probably due to the high reproductive rates of prokaryotes as well as their tendency to quickly follow environmental shifts: rapid abiotic recovery may thus shape and enable rapid prokaryotic recovery (Pressler et al., 2019; Whitman et al., 2019, 2022).

Fungal communities responded in a more complex and temporally variable manner. Immediately after the fire, general putative saprotrophs increased in relative abundance, which may be explained by the spike in fresh organic substrates such as lysed microbial cells and dead plant material, as well as by the nutrient pulse that favours more opportunistic fungi (Sun et al., 2015; Whitman et al., 2019; Certini et al., 2021). In addition, some of these positive responders are likely to be fire-adapted and particularly proliferate in immediate response to temperature and/or elevated pH (Raudabaugh et al., 2020; Arunrat et al., 2024). On the taxon level, for example, we identified members of the yeast- or mold-like Filobasidiaceae and Coniochaetaceae families as consistent and fast positive responders to fire (Fig. 3a). Particularly *Coniochaeta cipronana* obtained a high relative dominance. Members of *Coniochaeta* have been identified as pyrophilous, rapidly colonizing and saprotrophic fungi in other ecosystems (A. R. Nelson et al., 2022; Caiafa et al., 2023; Greenwood et al., 2023; Joukhajian et al., 2026). Other saprotrophic fungi that are considered pyrophilous are members of *Penicillium* (Aspergillaceae) (Mikita-Barbato et al., 2015; Whitman et al., 2019; Arunrat et al., 2024; Joukhajian et al., 2026), which interestingly had equal amounts of positive and negative responders in our study (Suppl. Fig. S5).

In contrast, more specialized litter- and wood-associated saprotrophic fungi tended to decline after burning, which we attribute to the fire-induced loss of litter as well as shrubs and trees. This explanation is further supported by the positive correlation between estimated litter cover and the relative abundance of putative litter saprotrophs in our plots (Spearman’s rho = 0.30, *P* = 0.004). Mycoparasites temporarily expanded, potentially capitalizing on an increased host abundance or on fungi rendered vulnerable by heat exposure (Semenova-Nelsen et al., 2019). Well-known mycorrhizal taxa seemed to primarily show negative responses: for example, arbuscular mycorrhizal fungi of the Glomeraceae family primarily decreased, and so did most members of the Helotiaceae and Hyaloscyphaceae families that contain many mycorrhizal fungi. This is corroborated by several studies that found negative effects on mycorrhizal fungi after fire across ecosystems, mostly driven by host mortality (Dove & Hart, 2017; Caiafa et al., 2023; Pulido-Chavez et al., 2023). In particular, we found a persistent decrease in the relative abundance of *Hyaloscypha variabilis* (Suppl. Fig. S5), an ericoid mycorrhizal fungi, which is in line with results from a burned down pine forest (Olchowik et al., 2021). These patterns of reduced mycorrhizal symbionts can possibly be explained by (i) direct mortality of the host plants and thus of their associated mycorrhizal fungi, and (ii) reduced reliance on symbioses under the temporary elevated nutrient availability, which may alter competitive balances in favour of saprotrophs (Treseder, 2004; Fox et al., 2022).

Most striking in our study was that, unlike prokaryotes, the timing and duration of fungal responses varied across taxa. Some fungi responded immediately, others showed delayed or short-lived responses, and few exhibited more persistent shifts, resulting in burned plots hosting a dynamically changing community composition over time, hence following a successional pathway. During the first two years of this post-fire fungal succession, we also observed a temporary decoupling between plants and fungi, as evidenced by the disruption of –otherwise strong– vegetation-fungal community correlations in burned plots (Fig. 5). Despite this delayed and slower recovery, fungal communities of burned plots did tend to converge towards intact plots at the end our study: in year three, differences in fungal community composition were only marginally significant (p = 0.053, Fig 1), and the positive correlation between vegetation- and fungal community composition had been restored (Fig 5f).

### 4.5 Conclusions

Our integrated three-year assessment of the effects of a flaming wildfire in a temperate peatland–heathland mosaic demonstrates that above- and belowground components recover at different rates. Soil physicochemical properties were only briefly affected: nutrient pulses of ammonium, nitrate, and bioavailable phosphorus in the year of the fire quickly returned to a baseline, reflecting the limited severity and surficial nature of the burn. Vegetation also recovered rapidly, with community composition largely converging with intact plots already within one year. In contrast, soil microbial communities, particularly fungi, displayed stronger, diverse and more persistent responses that followed a specific successional pathway. During the first two years post-fire, we witnessed a transient decoupling between plants and fungi in burned plots as evidenced by the temporary disruption of vegetation–fungal community correlations. In conclusion, our results highlight that swift aboveground recovery after a severe perturbation can mask prolonged belowground disruptions, with potential implications for decomposition, nutrient cycling, and plant–microbe interactions. By integrating vegetation, microbial, and soil dynamics, our study provides rare empirical evidence from an ecosystem without a natural fire regime and underscores the importance of considering multi-level ecosystem responses when assessing resilience and predicting the consequences of increasing wildfires under climate change.

## Acknowledgments

We gratefully acknowledge the assistance provided by Natuurpunt vzw, particularly Kris van der Steen and Guy Laurijssens. This study was financed by the Flemish Government (VR 2021 2602 DOC.0179/281S).

## Author contributions

KV, EV, RvD, KK & WJE designed and performed the field study; KV, EV, KK, YL, MC, SJ & WJE collected the data; LM analyzed the data; LM wrote the first draft of the manuscript and all authors contributed to the writing.

## Data availabilility

Abiotic data and vegetation relevees have been deposited in the Mendeley data repository (DOI: 10.17632/48nvyj2tk7.1). Raw sequences will be deposited in the SRA-NCBI database before publication.

## Supporting Information

The following Supporting Information is available for this article:

**Fig S1.**
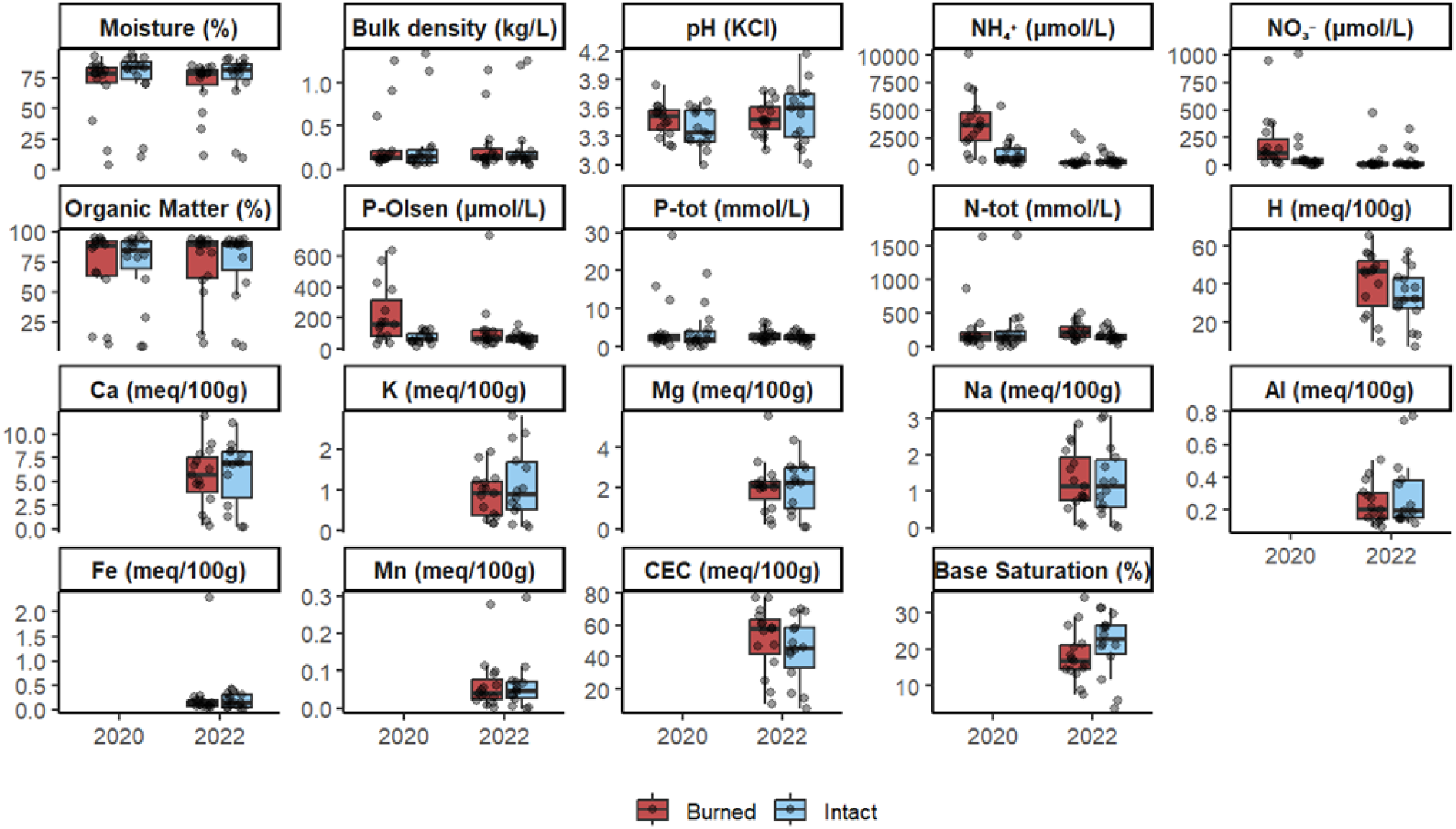
Soil physicochemical properties of paired burned (n=15) and intact (n=15) plots over the period 2020-2022

**Fig S2.**
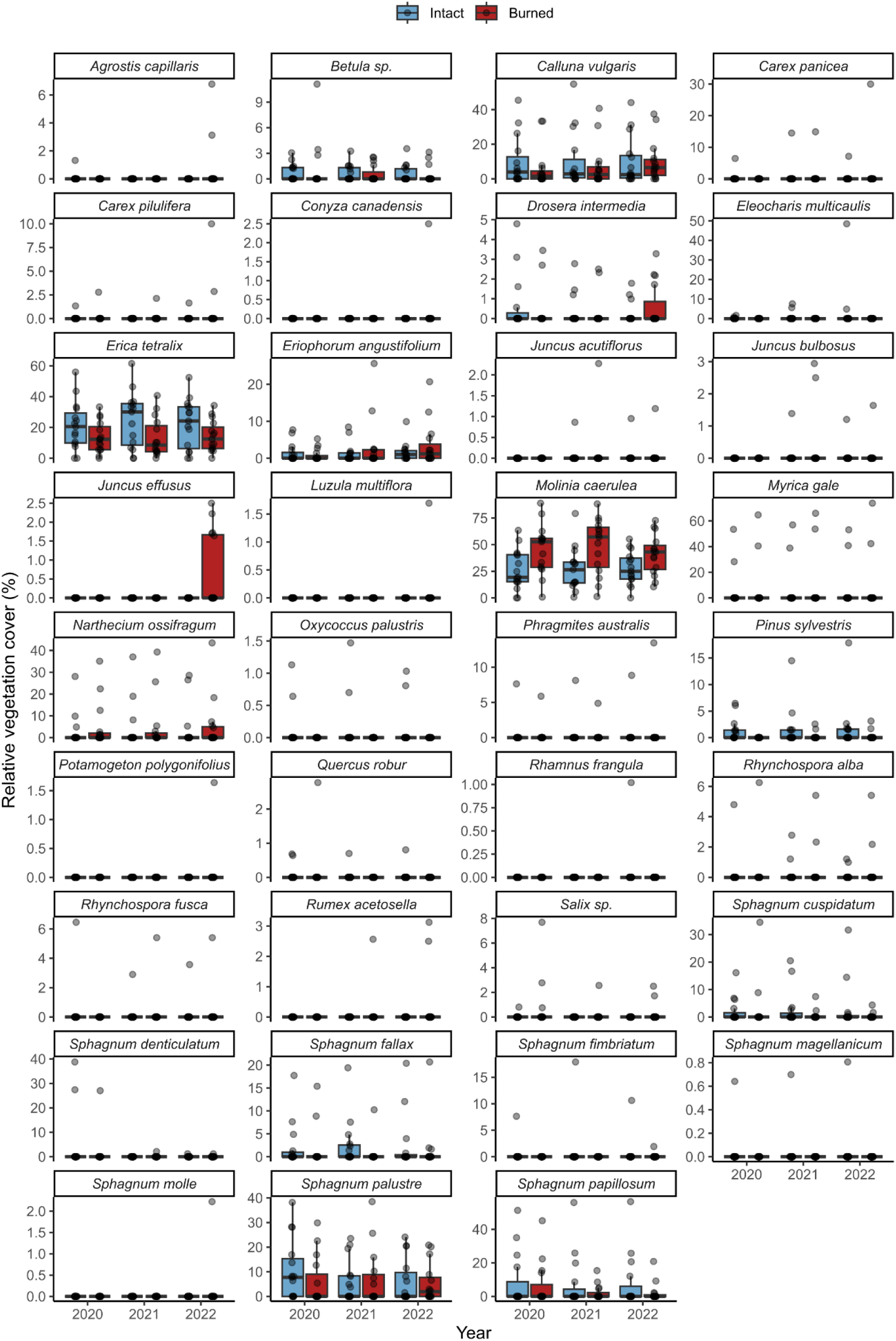
Relative plant species cover (%), calculated as individual species cover / total cumulative cover of all species (*100), in paired burned (n=15) and intact (n=15) plots over the period 2020-2022.

**Fig S3.**
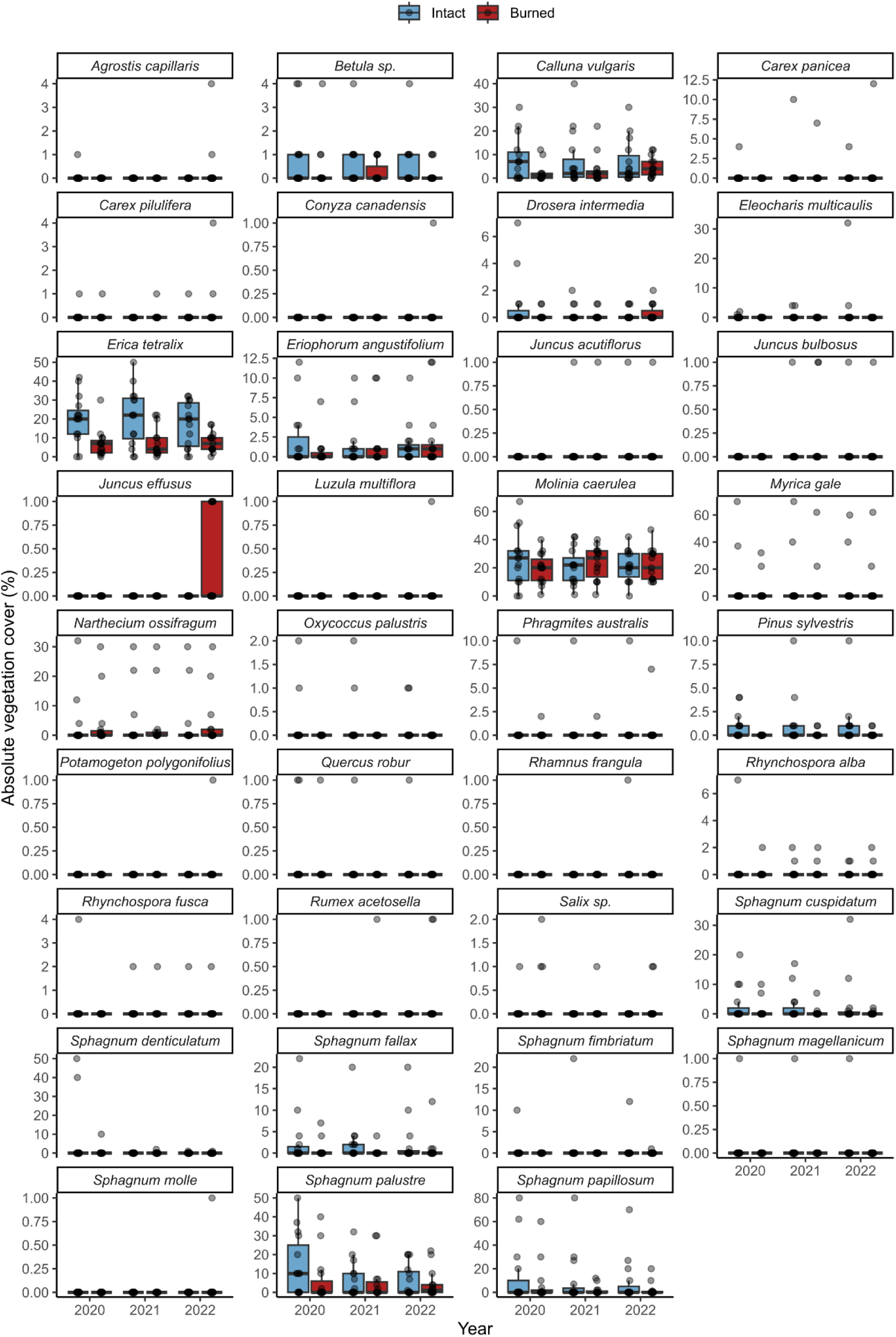
Absolute plant species cover (%), defined as the total cover per plot, in paired burned (n=15) and intact (n=15) plots over the period 2020-2022.

**Fig S4.**
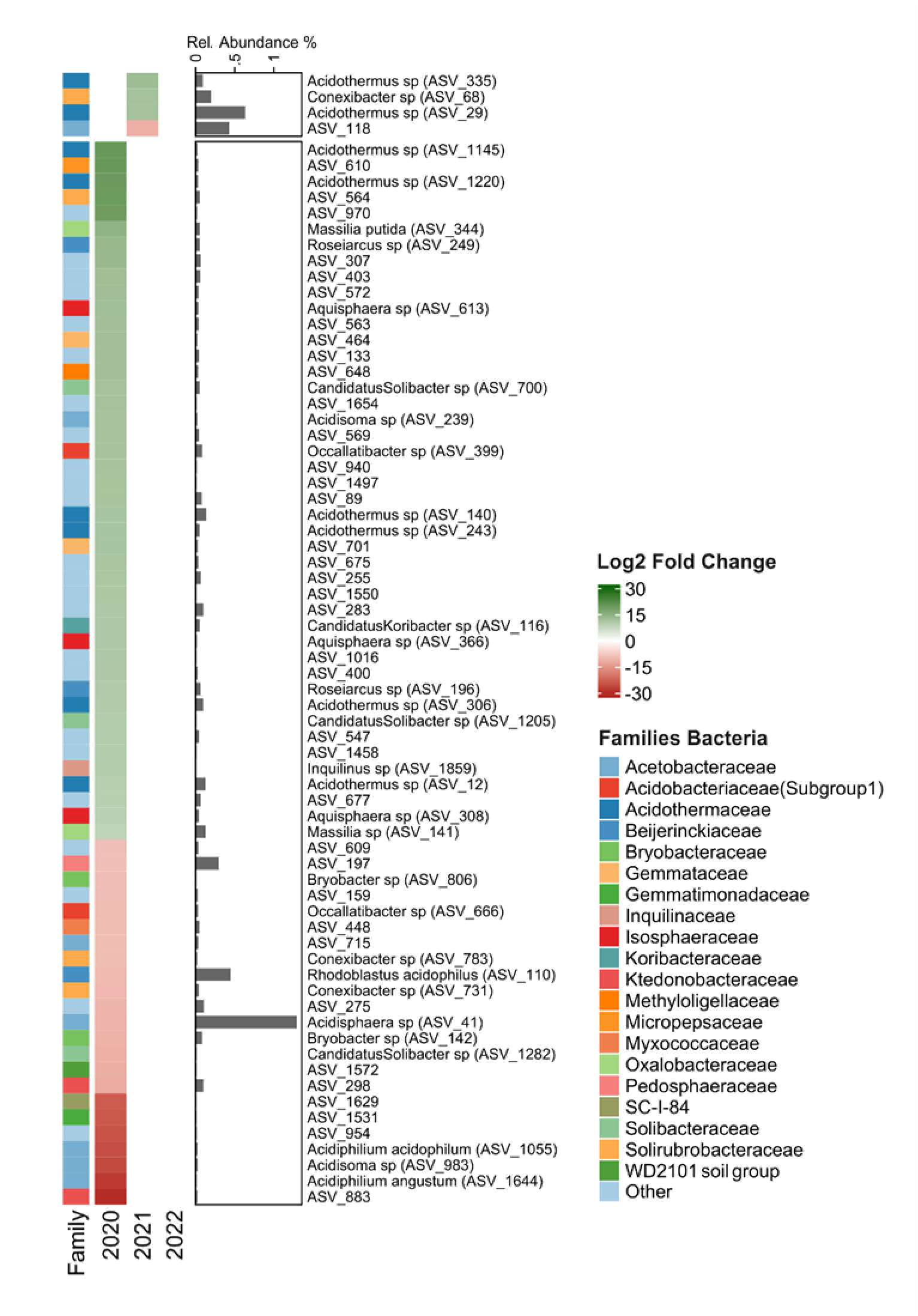
Responses of prokaryotic ASVs with a significant change in relative abundance (Log2 Fold change > |2|, p<0.05) following fire as identified by DESeq. This change can either be an increase (Log2FC > 2; green) or decrease (Log2FC < -2; red) in relative abundance. Responses are grouped by those that show a significant change in the first year (immediate) and those that only react in 2021 or 2022 (Delayed). Every row belongs to a single ASV with genus and species identification provided when available, family classification is given in the legend. Families with four or less responsive ASVs together with ASVs of unknown classifications are marked as ‘Other’. The column “Relative abundance” represents the average percentage of reads per ASV across all years and all plots combined.

**Fig S5.**
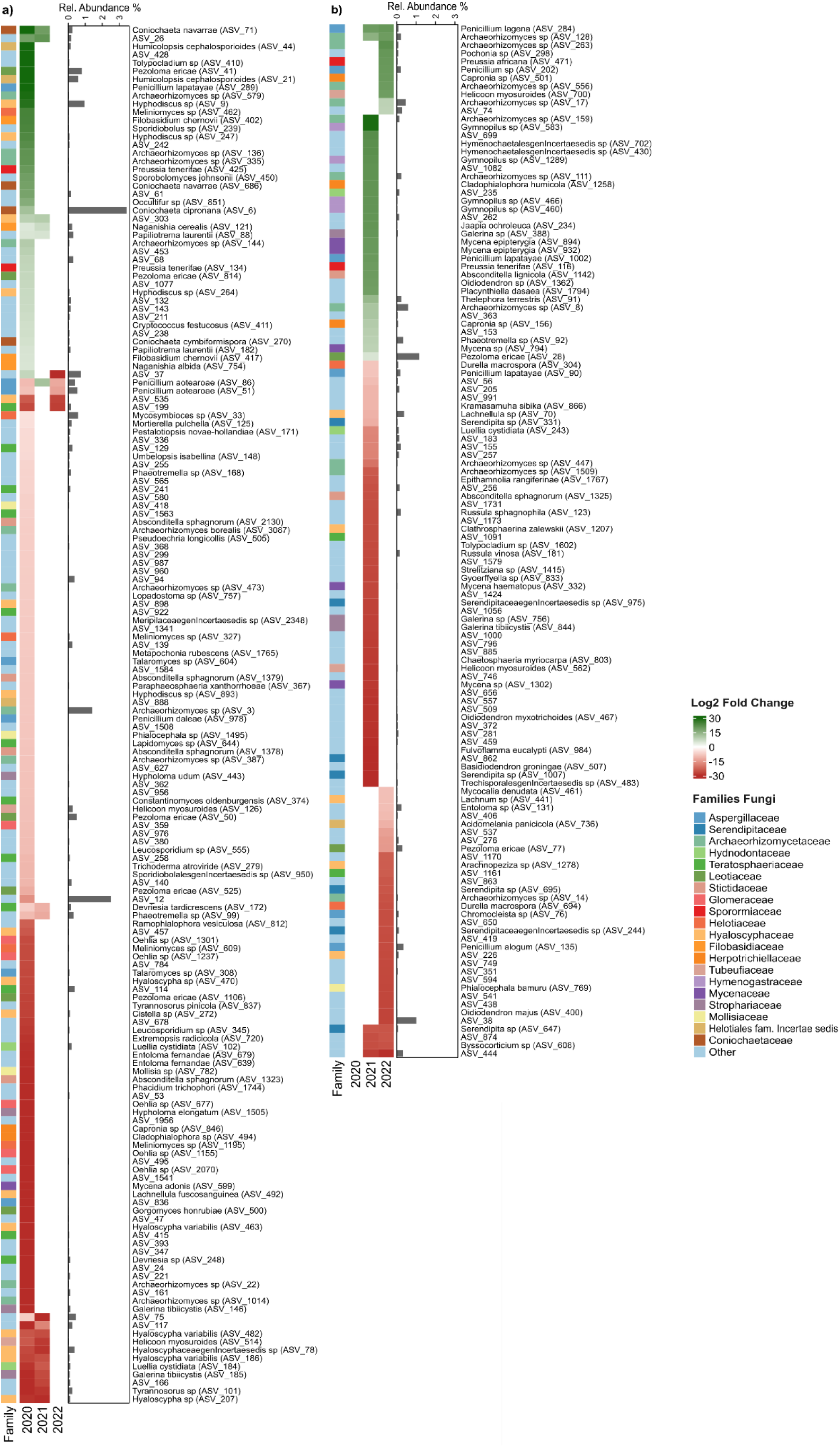
Responses of fungal ASVs with a significant change in relative abundance (Log2 Fold change > |2|, p<0.05) following fire as identified by DESeq. This change can either be an increase (Log2FC > 2; green) or decrease (Log2FC < -2; red) in relative abundance. Responses are grouped by those that show a significant change in the first year (immediate, a) and those that only react in 2021 or 2022 (Delayed, b). Every row belongs to a single ASV with genus and species identification provided when available, family classification is given in the legend. Families with four or less responsive ASVs together with ASVs of unknown classifications are marked as ‘Other’. The column “Relative abundance” represents the average percentage of reads per ASV across all years and all plots combined.

**Fig. S6.**
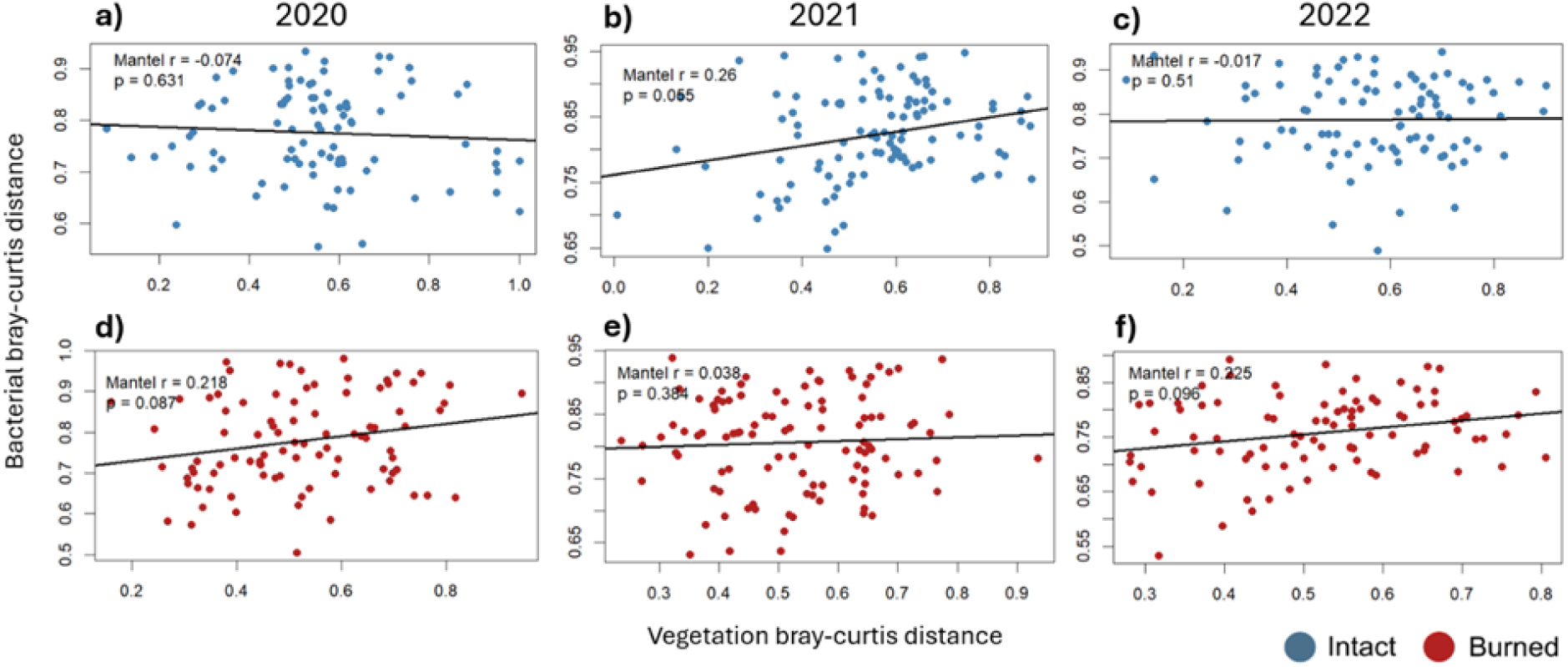
Correlations between prokaryotic and vegetation community composition in intact (panels a-c, n=15) and burned plots (panels d-f, n=15) for the period 2020-2022. Mantel’s correlation coefficients (rho) and p-values are indicated on the graphs.

